# The nuclear localization signal of CPSF6 governs post-nuclear import steps of HIV-1 infection

**DOI:** 10.1101/2024.06.20.599834

**Authors:** Nicholas Rohlfes, Rajalingam Radhakrishnan, Parmit K. Singh, Gregory J. Bedwell, Alan N. Engelman, Adarsh Dharan, Edward M. Campbell

## Abstract

The early stages of HIV-1 infection include the trafficking of the viral core into the nucleus of infected cells. However, much remains to be understood about how HIV-1 accomplishes nuclear import and the consequences of the import pathways utilized on nuclear events. The host factor cleavage and polyadenylation specificity factor 6 (CPSF6) assists HIV-1 nuclear localization and post-entry integration targeting. Here, we used a CPSF6 truncation mutant lacking a functional nuclear localization signal (NLS), CPSF6-358, and appended heterologous NLSs to rescue nuclear localization. We show that some, but not all, NLSs drive CPSF6-358 into the nucleus. Interestingly, we found that some nuclear localized CPSF6-NLS chimeras supported inefficient HIV-1 infection. We found that HIV-1 still enters the nucleus in these cell lines but fails to traffic to speckle-associated domains (SPADs). Additionally, we show that HIV-1 fails to efficiently integrate in these cell lines. Collectively, our results demonstrate that the NLS of CPSF6 facilitates steps of HIV-1 infection subsequent to nuclear import and additionally identify the ability of canonical NLS sequences to influence cargo localization in the nucleus following nuclear import.

**Author Summary:** During HIV-1 infection, the viral capsid, which encloses the viral genome and accessory proteins required for reverse transcription (RT) and integration, traffics towards the nucleus and enters through the nuclear pore complex (NPC). Following entry into the nucleus, RT is completed and viral capsid disassembles releasing the preintegration complex (PIC) to integrate with the host chromosome. In this study, we investigated the early HIV-1 host factor CPSF6, and specifically focused on the C-terminal short amino acid nuclear localization signal (NLS) in CPSF6, in mediating viral nuclear entry and subsequent gene expression. Altering the NLS in CPSF6 with NLS from other proteins, significantly impacted HIV-1’s ability to infect those cells. We further showed this defect in infection occurred at the level of viral integration. This study highlights the importance of the NLS in CPSF6 in dictating the NPC it associates with and its effect on HIV-1 infection. Moreover, our study emphasizes the function of NLS in targeting host cargos to different nuclear entry pathways.

## Introduction

A defining feature of lentiviruses, including human immunodeficiency virus-1 (HIV-1), is the ability to infect non-dividing cells, which requires import of the viral ribonucleoprotein complex into the target cell nucleus. This requires that the virus traverse nuclear pore complexes (NPCs) that exist within the nuclear membranes of non-dividing cells. NPCs are large, macromolecular complexes composed of multiple copies of about 30 nucleoporins (Nups) (1–3). Critically, NPCs are responsible for maintaining the nuclear permeability barrier, which restricts entry of many proteins while simultaneously allowing directed trafficking of hundreds to thousands of cargoes per second (4, 5). As a result, nucleocytoplasmic trafficking is a complex and tightly regulated process. To ensure the correct cargo is being transported into the nucleus, short amino acid sequences called nuclear localization signals (NLSs) are recognized by nuclear transport receptors to facilitate nuclear import (6, 7).

The HIV-1 capsid core is comprised of approximately 250 hexamers and exactly 12 pentamers of capsid protein (CA) forming a conical structure that houses the viral genome and essential enzymes (8–12). CA is the primary viral determinant driving nuclear import (13), and CA accordingly directly interacts with several Nups including Nup358 (14–16), Nup153 (15, 17–22), Nup58 (15, 21), Nup98 (15, 21), and Nup62 (15, 20), as well as with CPSF6 (15, 17, 19, 21, 23, 24). Following trafficking through the cytoplasm, the core docks at the cytoplasmic face of the NPC via interactions with Nup358 (16, 25). This promotes passage through the NPC (16, 25, 26), leading to docking interactions with Nup153 on the nucleoplasmic side of the NPC (20), where interactions with CPSF6 are thought to release the core into the nucleus (27, 28). CPSF6 traffics with the core further into the nucleus to subnuclear bodies called nuclear speckles (NSs) (27, 29–31). NS are found near gene-dense regions of chromatin and prior work has shown that HIV-1 preferentially targets these regions for proviral integration (30, 32). Importantly, HIV-1 capsid mutants defective for CPSF6 binding display aberrant integration site selection, highlighting the impact of CPSF6 for integration (16, 33).

CPSF6 contains a functional NLS at its C-terminus, and as a result is predominantly found in the nucleus of cells (34–36). CPSF6 is expressed as two splice isoforms. Isoform 1, the major form, is composed of 551 residues while isoform 2, containing 588 amino acids, carries internal residues encoded by exon 6 that are removed from isoform 1 by alternative splicing (34). A truncation of CPSF6 isoform 2 containing amino acids 1-358 (CPSF6-358) was identified by the KewalRamani group, where they noted the protein was mislocalized to the cytoplasm and acted as an artificial restriction factor (34). This finding led to identification of the N74D capsid mutant, which is defective in its ability to bind to CPSF6 (34). In the same study, using siRNA knockdown of various Nups, they identified that the N74D CA mutant and wildtype (WT) CA showed variable sensitivities to Nup knockdown, highlighting their utilization of different nuclear import pathways (34). Investigation of the P90A capsid mutant also found that it utilizes a nuclear import pathway distinct from WT virus (16, 37). Our previous studies using a nuclear pore blockade have also supported the idea of multiple nuclear import pathways (38). We observed that infection mediated by WT CA was potently inhibited by Nup62 blockade, but infection mediated by N74D and P90A CA was comparatively less sensitive to insensitive to the Nup62-mediated NPC blockade, suggesting the utilization of distinct nuclear import pathways, likely utilizing different nuclear pore complexes (38). Recently, several groups reported how swapping an NLS can alter sensitivity to the restriction factor MX2 (39, 40). Some NLSs retained the restriction phenotype, whereas others rendered HIV-1 less susceptible to MX2 restriction (39, 40). These findings showed that some, but not all, import pathways retained MX2 restriction activity and that the NLS determined which pathway was used.

To better understand the role of CPSF6’s NLS, we attached heterologous NLSs to CPSF6-358 to determine the impact of divergent NLSs on nuclear import and infection by HIV-1 and related lentiviruses. We observed that the addition of NLS sequences that restored the nuclear localization of CPSF6 also promoted the nuclear import of HIV-1 during infection. However, despite effective nuclear import, we noted that some NLSs promoted a dysfunctional form of nuclear import that did not facilitate subsequent intranuclear steps of infection, including integration. These results demonstrate that the NLS promoting the nuclear import of HIV-1 can impact the nuclear stages of infection. Previous reports have shown that CPSF6’s C-terminal intrinsically disordered region (IDR), which is the protein’s NLS (39), confers NS colocalization and condensation to heterologous fusion partners such as green fluorescent protein (GFP) (35, 41, 42). Our results support these findings, as we found only the full-length CPSF6, which contains an intact IDR, formed condensates following HIV-1 infection. However, our findings show re-localization to NSs can occur independently of the IDR, as CPSF6-NLS constructs containing the SV40 or C-MYC NLS effectively trafficked HIV-1 to NSs despite lacking the IDR. These data show that specific NLSs license subsequent trafficking and integration by lentiviruses and further refines our understanding of the nuclear import pathways utilized by HIV-1. These data also provide evidence that NLSs possess the ability to influence post-nuclear import activities of their cargoes.

## Results

### Heterologous NLSs can confer CPSF6-358 nuclear localization

To understand more about CPSF6’s NLS and the influences it has on post-nuclear HIV-1 biology, we used the well described truncation mutant containing amino acids 1-358 (CPSF6-358), which lacks the C-terminal IDR, also called an RS-like domain (RSLD) (34). The RSLD contains the functional NLS for CPSF6, and as a result, CPSF6-358 displays a significant cytoplasmic localization, in contrast to full-length CPSF6, which is predominantly nuclear (36). We then introduced various NLS’s on the C-terminus of CPSF6-358 to rescue nuclear localization through several distinct nuclear import pathways (Fig 1A and Table 1). We chose representative NLSs, which were either cellular or viral in origin, and spanned across the different NLS classifications. To understand the impact of these NLSs on infection, we first knocked out endogenous CPSF6 in HeLa cells and generated stable cell lines expressing the indicated CPSF6-NLS chimeras (Fig 1B-C). We observed that several of the NLSs (SV40, C-MYC, NP, Rac3, HNRNP K, HNRNP A1, and MX2) effectively rescued nuclear localization of the CSPF6-358 construct (Fig 1D, Table 1). Notably, some NLSs (DDX21, RB, HTLV-1 Rex, and UL79) were unable to confer effective nuclear localization, and the corresponding CSPF6-358 chimeric proteins retained cytoplasmic distribution (Fig 1D-E, Table 1). These results are in line with previous studies of MX2 localization, which showed variable degrees of nuclear localization of heterologous MX2 constructs across the tested NLSs (39, 40). Overall, some, but not all, of our chimeric CPSF6-NLS constructs have restored nuclear localization.

**Fig 1.**
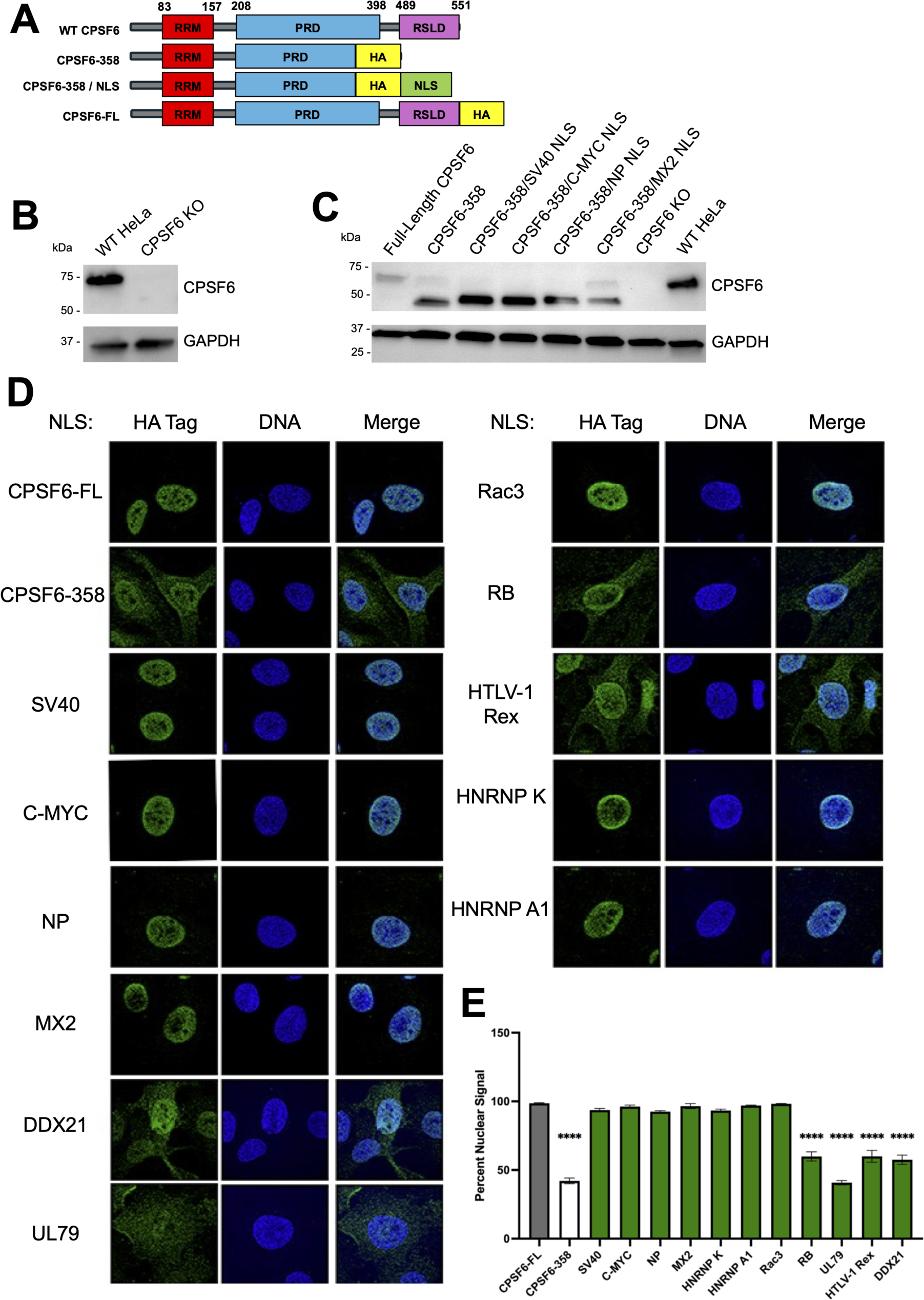
Heterologous NLSs can confer CPSF6-358 nuclear localization. (A) Schematic design of chimeric CPSF6-NLS constructs showing NLSs being fused to the C-terminus of CPSF6-358 derived from human isoform 1. (B) Western blot analysis of HeLa cells using anti-CPSF6 antibody to confirm CRISPR/Cas9-mediated knockout of endogenous CPSF6. (C) Western blot analysis of stably transduced HeLa cells using anti-CPSF6 antibody to detect CPSF6-NLS construct expression following 48 h doxycycline induction. Anti-GAPDH antibody used as loading control. (D) Immunofluorescent microscopic images stained using anti-HA (green) and Hoechst (blue) antibodies to visualize CPSF6-NLS construct localization. Representative of 3 independent experiments with at least 10 images per experiment. (E) Percent nuclear signal was determined using ImageJ. Statistical analysis in E was determined using one-way ANOVA. Significant differences are indicated: **** P<0.0001. RRM – RNA Recognition Motif. PRD – Proline Rich Domain. RSLD – Arginine Serine-Like Domain. HA – Hemagglutinin tag. NLS – Nuclear Localization Signal. CPSF6-FL – Full Length CPSF6.

**Table 1.** Nuclear localization signals fused to CPSF6-358 and their effect on construct localization and WT HIV-1 gene expression.

| NLS | NLS Type | Construct Localization<sup>a</sup> | Percent Infectivity<sup>b</sup> | Reference |
| --- | --- | --- | --- | --- |
| *CPSF6* | RS-NLS | Nuclear | 100 ± 17 |  |
| *CPSF6-KO* | - | - | 107 ± 22 |  |
| *CPSF6-358* | None | Cytoplasmic | 10 ± 8 |  |
| *SV40* | Monopartite Class 1 | Nuclear | 83 ± 32 | (43) |
| *C-MYC* | Monopartite Class 2 | Nuclear | 141 ± 61 | (44) |
| *DDX21* | Monopartite Class 3 | Cytoplasmic | 5 ± 1 | (44) |
| *NP* | Bipartite | Nuclear | 32 ± 9 | (43) |
| *RB* | Bipartite | Cytoplasmic | 15 ± 17 | (45) |
| *Rac3* | Bipartite | Nuclear | 140 ± 45 | (45) |
| *UL79* | PY-NLS | Cytoplasmic | 11 ± 12 | (46) |
| *HNRNP A1* | PY-NLS | Nuclear | 70 ± 17 | (47) |
| *HNRNP K* | n/a | Nuclear | 80 ± 18 | (48) |
| *HTLV-1 Rex* | n/a | Cytoplasmic | 7 ± 2 | (49) |
| *MX2* | n/a | Nuclear | 14 ± 10 | (50) |

<sup>a</sup>Constructs retaining cytoplasmic distribution shown in red.
<sup>b</sup>Percent infectivity (mean ± SEM) from n ≥ 3 independent experiments, with each experiment containing at least technical duplicates. Constructs resulting in significant infectivity reductions shown in red.

### Nuclear localization of CPSF6-358 does not uniformly rescue HIV-1 infection

CPSF6 is an HIV-1 host factor and binds directly to the HIV-1 capsid to aid in nuclear entry and integration site targeting (17, 23, 24, 34). To further understand if changing CPSF6’s NLS impacts HIV-1 infection, we utilized single-round HIV-1 luciferase reporter virus to infect our chimeric CPSF6 cell lines. To highlight phenotypes associated with NLS functionality, preliminary experiments were conducted with growth-arrested cells. Consistent with previous studies (34), CPSF6-KO did not perturb HIV-1 infectivity in HeLa cells, and expression of CSPF6-358 in CPSF6-KO cells potently inhibited infection (Fig 2A). A similar inhibition was observed in cells expressing chimeric NLSs that failed to confer CPSF6-358 nuclear localization (DDX21, RB, HTLV-1 Rex, and UL79) (Fig 2A, striped bars). However, we observed three distinct phenotypes amongst cells expressing CPSF6 chimeras with restored nuclear localization. We found that the C-MYC and Rac3 NLS not only fully rescued infection, but resulted in a small but significant increase in infectivity compared to the infection observed in CPSF6-KO and CPSF6-FL cells (Fig 2A). In contrast, the SV40, HNRNP K, and HNRNP A1 NLSs rescued infection to levels that were indistinguishable from CPSF6-FL (Fig 2A). Finally, HIV-1 infection was significantly reduced in cells expressing the CSPF6 chimeras with NP and MX2 NLSs (Fig 2A), despite these NLSs conferring effective CPSF6-358 nuclear localization. These phenotypes persisted across several viral titrations, showing that the differences in infectivity were concentration independent (Fig 2B). We observed similar infectivity findings using a fluorescent reporter virus, revealing that the results were independent of the utilized viral reporter gene (Fig 2C). Additionally, these results were independent of cell cycle arrest, as we observed similar phenotypes when cells were not growth arrested with aphidicolin (Fig 2D). These results show that simply conferring constitutive CPSF6-358 nuclear localization is insufficient to restore HIV-1 infectivity and is impacted by the NLS used to mediate CPSF6-358 nuclear localization.

**Fig 2.**
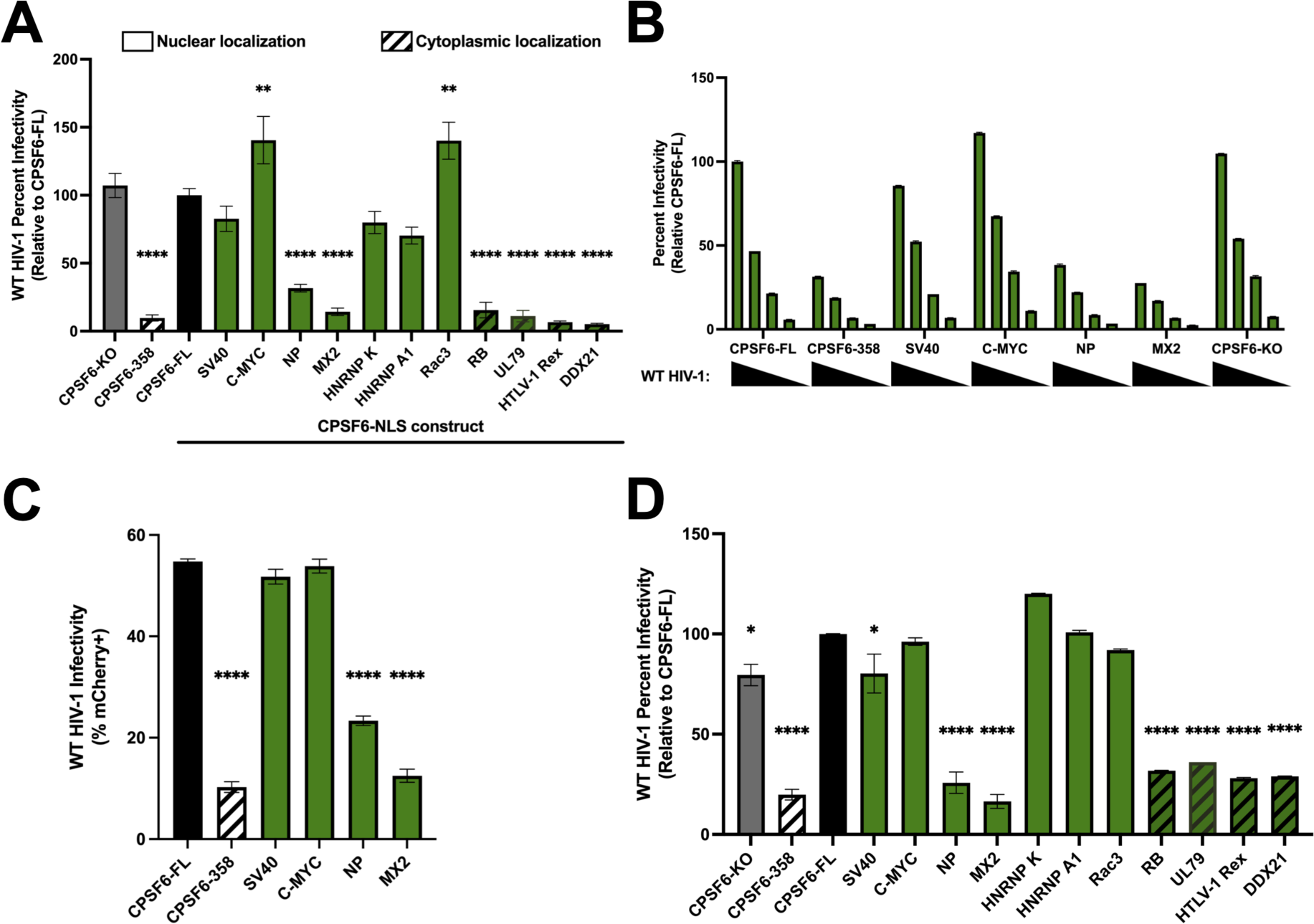
Nuclear localization of CPSF6-358 does not uniformly rescue HIV-1 infection. (A) WT HIV-1 firefly luciferase reporter virus infectivity was measured 48 h post-infection in growth arrested HeLa cells. Infectivity was normalized to cells expressing full-length CPSF6. Striped bars represent constructs with cytoplasmic fractions. (B) Serial dilutions show that infectivity phenotypes hold across several viral concentrations in growth arrested cells. (C) WT HIV-1 mCherry reporter virus was used to infect growth arrested HeLa cell lines and infectivity was determined using flow cytometry. (D) Dividing HeLa cells infected with WT HIV-1 firefly luciferase reporter virus displays comparable phenotypes to non-dividing cells. Results (mean ± SEM) are representative from at least 3 independent experiments with at least technical duplicates. Statistical analysis in A was determined using one-way ANOVA. Significant differences are indicated: ** P<0.01, **** P<0.0001. CPSF6-FL – Full Length CPSF6.

The ability to influence infection positively or negatively was not dependent on the type of NLS, which can be categorized based on their sequence and the nuclear transport machinery that recognizes the NLS (51). RB, Rac3, and NP are three examples of bipartite NLSs that conferred three distinct phenotypes under our assay conditions (Table 1). Rac3 and NP conferred CPSF6-358 nuclear localization, whereas the RB NLS chimera remained cytoplasmic. Additionally, Rac3 fully rescued infectivity while NP resulted in an infectivity defect. Additionally, we tested two PY-NLS’s, UL79 and HNRNP A1, and found that HNRNP A1 conferred CPSF6-358 nuclear localization and supported HIV-1 infection similarly as CPSF6-FL, while UL79 failed to rescue both CPSF6-358 nuclear localization and HIV-1 infection (Table 1). These findings suggest that the ability to influence HIV-1 infection cannot be explained by NLS classification. Together, these results display the importance of the nuclear import pathway utilized by CPSF6 for productive HIV-1 infection.

### CPSF6-NLS chimeras show virus-specific influences on infection

To ensure that the observed changes in viral gene expression were driven through CA-CPSF6 interactions and not a deleterious effect of chimeric protein expression on cell function, we infected these cells with the CPSF6 binding deficient HIV-1 CA mutant N74D (34). This mutant contains a point mutation in the CA-CPSF6 binding pocket, rendering CA unable to bind to CPSF6. Importantly, this CA mutant retains the ability to productively infect HeLa cells (34). When we infected our chimeric cell lines with N74D virus, we observed no differences in viral gene expression, demonstrating that the differences in WT HIV-1 infectivity observed in cells expressing CPSF6 chimeras is dependent of CA binding (Fig 3A).

**Fig 3.**
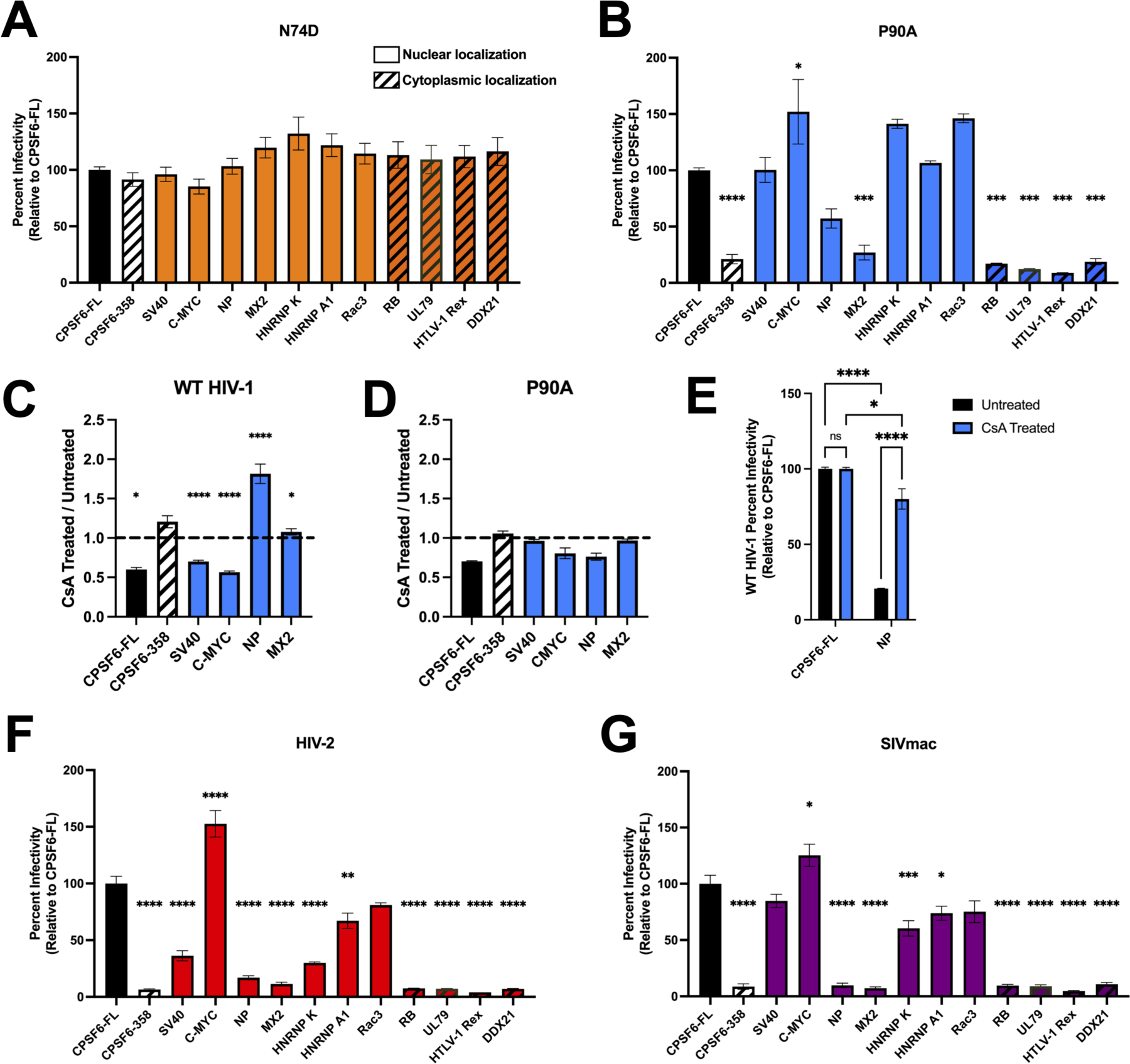
CPSF6-NLS chimeras confer virus-specific infectivity defects. (A) Infection with HIV-1 CA mutant N74D luciferase-reporter virus in growth arrested HeLa cells reveals a dependence on CA-CPSF6 binding for the observed changes in WT HIV-1 infectivity. (B) Infection with P90A CA mutant in growth arrested HeLa cells results in less-severe infectivity defect with CPSF6-NP NLS chimera cells, while CPSF6-HNRNP K NLS cells trend to be more sensitive to infection relative to WT virus. (C) Relative infectivity in growth of CsA-treated growth arrested cells to untreated HIV-1-infected growth arrested cells displays a decrease to no change in infectivity for all CPSF6-NLS cell lines except for CPSF6-NP NLS cells, which display an almost twofold increase in sensitivity to infection. (D) Comparison CsA treated vs untreated P90A infected cell lines displays little to no change across all growth arrested cell lines. (E) CsA treatment of WT HIV-1 infected CPSF6-FL and CPSF6-NP NLS cell lines following growth arrest displays significant increases in infectivity in CPSF6-NP NLS cells following CsA treatment. Infection with non-HIV-1 primate lentiviruses reporters, HIV-2 (E) and SIVmac (F) in growth arrested cells. Relative to HIV-1 infection, there are several decreases in HIV-2 infectivity, whereas SIVmac appears fairly similar to HIV-1. Results (mean ± SEM) are representative from 3 independent experiments with at least technical duplicates. Statistical analysis in A,B,F,G was determined using one-way ANOVA and C-E was determined using two-way ANOVA. Significant differences are indicated: * P<0.05, ** P<0.01, *** P<0.001, **** P<0.0001. CsA – Cyclosporine A. CPSF6-FL – Full Length CPSF6.

The P90A HIV-1 CA mutant is defective in its ability to bind to cyclophilin A (52–54), which is another host factor implicated in HIV-1 nuclear import (55). Importantly, P90A retains the ability to bind to CPSF6, but has been shown to utilize a nuclear import pathway distinct from WT HIV-1 (16). To further explore the role of the CA-CPSF6 interaction in HIV-1 infection, we infected our cell lines with the P90A mutant virus. We observed defective infection for cells expressing the cytoplasmic localizing CPSF6-NLS mutants, while cells expressing the NP and MX2 NLS variants again supported reduced levels of HIV-1 infection (Fig 3B). Cells expressing SV40, C-MYC, HNRNP K, HNRNP A1 and Rac3 chimeras supported P90A infection to a similar degree as observed for WT HIV-1 (Fig 3B). However, we noticed that the reduction in infectivity of the P90A virus was more modest in cells expressing the CPSF6-NP NLS than was observed in cells expressing the CPSF6-MX2 NLS. To corroborate this observed difference, we infected our cell lines with WT HIV-1 in the presence of cyclosporin A (CsA). CsA blocks cyclophilin A binding to CA and thus displays a phenotype similar to the P90A mutation (52–54). We found that CsA treatment of WT HIV-1 infected CPSF6-NP chimeric cell lines resulted in significantly increased infectivity relative to untreated cells, while other chimeric CPSF6-NLS cell lines displayed no change or decreased infectivity following CsA treatment (Fig 3C, E). Notably, CsA treatment of P90A infected cells resulted in little to no effect on infection in all cell lines tested, confirming that the observed changes were specific to the CA-cyclophilin interaction and not off-target CsA effects (Fig 3D). This demonstrates that disruption of CA-cyclophilin binding can modulate the ability of some CPSF6-NLS chimeras productively support HIV-1 infection.

Binding to CPSF6 is a shared characteristic amongst primate lentiviruses (32). We therefore wondered how CPSF6-NLS chimera expression would impact infection with HIV-2 and simian immunodeficiency virus from macaque (SIVmac). Interestingly, we found that CPSF6-SV40 NLS and CPSF6-HNRNP K NLS conferred strong HIV-2 infectivity defects, with CPSF6-HNRNP A1 conferring a more modest infection defect (Fig 3D), as compared to the effects these three proteins had on HIV-1 infection (Fig 2A). Akin to the HIV-1 results, HIV-2 displayed a strong increase in infectivity in CPSF6-C-MYC NLS expressing cells (Fig 3D). SIVmac displayed similar phenotypes to HIV-1, with the exception that SIVmac infection was modestly reduced in HNRNP K NLS expressing cells (Fig 3E). These results show that while the ability of CPSF6-NLS constructs to facilitate infection, was generally conserved among viruses with capsids that bind CPSF6, there are some differences in the way P90A and other primate lentirivurses can utilize these constructs to support productive infection.

### CPSF6-NLS chimeras display cell-type specific phenotypes

To test the CPSF6-NLS chimera constructs in more physiologically relevant settings, we depleted endogenous CPSF6 from T cell (SupT1, Jurkat) and monocyte-derived macrophage (THP-1) cell lines and generated stable cells expressing four representative CPSF6-NLS chimeras that were mapped in HeLa cells as constitutively nuclear (S1A-C Fig). In these cells, the CPSF6-SV40 NLS chimera rescued ∼50% of T cell infection (Fig 4A, B), whereas it fully rescued infection of THP-1 cells (Fig 4C). Additionally, CPSF6-C-MYC NLS expression increased THP-1 infectivity to about 200% relative to full length CPSF6 (Fig 4C), but rescued T cell infectivity to about 100% (Fig 4A, B). CPSF6-NP NLS and MX2 NLS constructs in T cells and macrophages behaved largely similarly as they did in HeLa cells, with the exception that the CPSF6-NP NLS chimera supported comparatively less infection of THP-1 cells (Fig 4A, B). Together, these results reveal the target cell can impact the ability of HIV-1 to utilize chimeric CPSF6-NLS constructs during infection.

**Fig 4.**
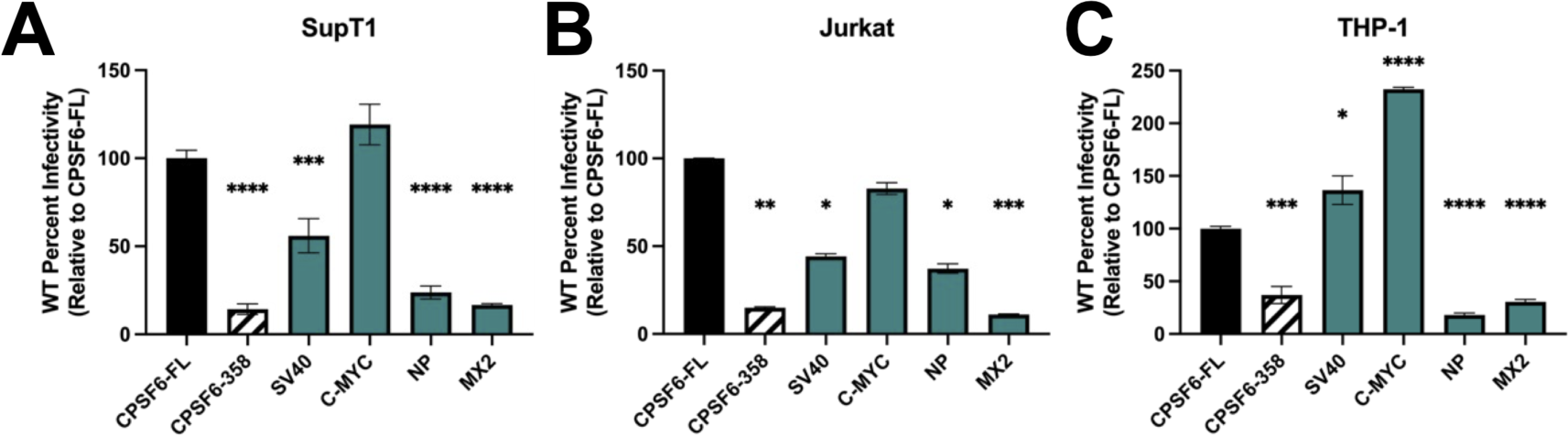
CPSF6-NLS chimeras display cell-type specific phenotypes. CPSF6-KO SupT1 (A) and Jurkat (B) T-cells and THP-1 monocytes were stably transduced with dox-inducible CPSF6-NLS constructs. T cells were growth arrested prior to infection. WT HIV-1 infection was determined using luciferase assay. Notably, T cells expressing CPSF6-SV40 NLS were significantly less infectable than corresponding HeLa cell lines. (C) THP-1 cell lines were differentiated to macrophages and infected with WT HIV-1. Notably, CPSF6-SV40 NLS THP-1 cells supported greater levels of infection than analogous T cell lines, whereas CPSF6-C-MYC NLS strongly increased infectivity relative to CPSF6-FL. Results (mean ± SEM) are representative from 3 independent experiments with at least technical duplicates. Statistical analysis was determined using one-way ANOVA. Significant differences are indicated: * P<0.05, ** P<0.01, *** P<0.001, **** P<0.0001.

### Heterologous NLSs significantly impact intranuclear stages of HIV-1 infection

Due to the CPSF6-NLS chimera constructs restoring nuclear localization of CPSF6-358 (Fig 1C), we next asked if defective HIV-1 gene expression was a result of defective nuclear import in these cells. Nuclear import is commonly quantified by measuring the accumulation of 2-long terminal repeat (LTR)-containing circles, which are a byproduct of the unintegrated HIV-1 genome being circularized by the host non-homologous end-joining machinery (56). We infected the chimeric NLS HeLa cell lines with WT HIV-1 and measured the formation of total HIV-1 DNA (late reverse transcription products) and 2-LTR circles by qPCR. All tested chimeric NLS cell lines supported similar levels of HIV-1 DNA synthesis (Fig 5A). CPSF6-358 expression, relative to full-length CPSF6, significantly reduced 2-LTR circle formation (Fig 5B), consistent with previous studies (34, 36). Interestingly, the CPSF6-NLS chimeric cell lines supported similar levels of 2-LTR circle accumulation (Fig 5B), despite the reduced levels of infectivity observed in CPSF6-358 NP NLS and MX2 NLS-expressing cells (Fig 2A). To further examine the ability of these constructs to mediate nuclear import, we utilized fluorescently tagged Gag-Integrase Ruby (GIR) virus to track the movement of individual virus particles (Fig 5C, E). Using IMARIS software, we formed a mask around the Hoechst signal and determined the percentage of our labeled particles within the nucleus. Consistent with our 2-LTR circle measurements, the CPSF6-NLS chimeric cell lines supported similar levels of GIR virus nuclear localization (Fig 5C, E). These results indicate that HIV-1 efficiently enters into the nuclei of our chimeric CPSF6-NLS cell lines and suggest a post-nuclear entry defect in the subset of these cells that results in significant reductions of WT virus infection.

**Fig 5.**
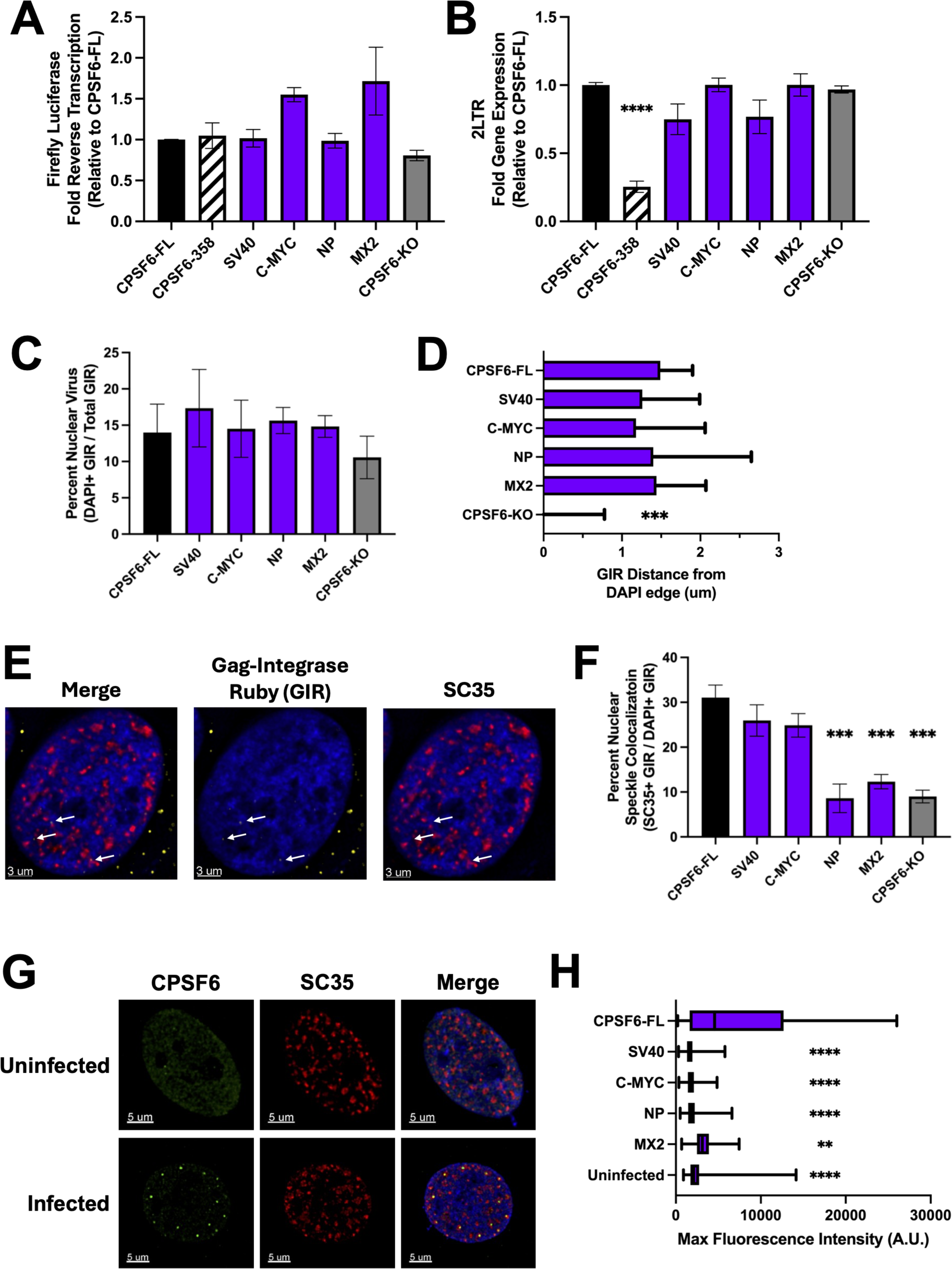
NLS on CPSF6 significantly impacts intranuclear stages of HIV-1 infection.

(A) Late reverse transcription was measured via firefly luciferase from the viral genome. (B) 2-LTR circles were measured and normalized to CPSF6-FL results. qPCR analysis was performed on genomic DNA isolated from WT HIV-1 infected growth arrested HeLa cells. (C) Gag-Integrase Ruby (GIR) labeled HIV-1 viral particles were used to infect growth arrested HeLa cell lines. Nuclear virus was determined using IMARIS software and creating a mask around the nuclear Hoechst stain, then determining percent of total GIR particles within the mask. There were no observable differences in percentage of nuclear GIR across the indicated cell lines. (D) Nuclear penetration was determined via IMARIS software, measuring Hoechst-positive GIR distance from Hoechst edge. GIR in CPSF6-KO cells remained nuclear edge proximal, whereas other CPSF6 chimeras facilitated nuclear invasion. (E) Representative images of GIR infected HeLa cells, with Hoechst (blue), GIR (yellow), and SC35 (red). (F) NS colocalization was determined via GIR infection and subsequent SC35 staining. Percentage of nuclear GIR colocalizing with SC35 was determined using IMARIS software. HIV-1 in CPSF6-NP NLS, CPSF6-MX2 NLS, and CPSF6-KO cells displayed decreased NS colocalization. (G) Representative images showing CPSF6-FL distribution in uninfected versus HIV-1 infected cells, with Hoechst (blue), CPSF6 (green), and SC35 (red). (H) CPSF6 condensation was determined via HA-tag staining at 18 h post-infection. Masks were made around SC35 stain using IMARIS and fluorescent intensity of CPSF6 construct stain within SC35 masks was determined. CPSF6-FL supported much brighter spots consistent with condensation, whereas CPSF6 distribution remained comparable to uninfected cells across other cell lines. All imaging experiments were performed on growth arrested cells. Results in A-C and F (mean ± SEM) are representative from 3 independent experiments with at least technical duplicates. Results in D show median+95% CI. Results in H show box and whiskers from min to max. At least 10 images were taken for each condition across 3 independent experiments. Statistical analysis in A-C and F was determined using one-way ANOVA and in D and H using Kruskal-Wallis test. Significant differences are indicated: * P<0.05, ** P<0.01, *** P<0.001, **** P<0.0001. GIR – Gag-Integrase Ruby. CPSF6-FL – Full-length CPSF6.

Following HIV-1 nuclear import, HIV-1 traffics away from the nuclear envelope and further into the nucleus (27, 57). The trafficking away from the nuclear envelope is licensed by CPSF6, as CPSF6-KO cells display increased nuclear envelop adjacent virus (27, 57). We quantified viral nuclear penetration by measuring GIR distance to the nearest Hoechst edge at 8 h post-infection. As expected, HIV-1 in CPSF6-KO cells remained near the nuclear periphery (Fig 5D). Viruses in CPSF6-NLS chimera expressing cells, however, penetrated deeper into the nucleus, comparably to cells expressing full-length CPSF6 (Fig 5D). The CA-CPSF6 interaction facilitates viral intranuclear trafficking to NSs (29, 30, 32, 58, 59). To determine if chimeric CPSF6-NLS constructs impair HIV-1 intranuclear trafficking to NSs, we utilized GIR virus in combination with an SC35 stain to visualize colocalization with NSs at 8 h post-infection (Fig 5E). CPSF6-KO cells displayed a reduction in GIR colocalization with NSs, as described previously (29–31, 58). However, the SV40 and C-MYC NLS constructs effectively facilitated localization to NSs (Fig 5F). Alternatively, the NP and MX2 NLS constructs conferred significant decreases in HIV-1 colocalization with NSs (Fig 5F). During infection, CA induces the formation of CPSF6 biomolecular condensates, which colocalize with NSs (58–60). Due to the SV40-NLS and C-MYC NLS chimeras facilitating viral localization to NSs, we wondered if these mutants were inducing the formation of biomolecular condensates, as previously described for full-length CPSF6 (30, 58–60). We measured CPSF6 signal intensity colocalizing with SC35 and found that only the full-length CPSF6 condensed with NSs, whereas the CPSF6-NLS chimeras retained diffuse nuclear localization similar to uninfected cells (Fig 5G, H). Taken together, our results show that a subset of our CPSF6-NLS constructs that supported maximal or partial HIV-1 infection nevertheless supported equivalent levels of HIV-1 nuclear entry. Moreover, following nuclear entry, the SV40 and C-MYC NLS were able to facilitate HIV-1 nuclear trafficking and NS localization despite a lack of biomolecular condensation. Conversely, the NP and MX2 constructs, despite facilitating nuclear penetration, were unable to facilitate NS localization or biomolecular condensation of CPSF6.

### CPSF6-NP and MX2 NLS chimeras reduce HIV-1 integration and influence integration site selection

The substrate for HIV-1 integration is the linear viral DNA made by reverse transcription. HIV-1 integration favors active chromatin including DNA in close proximity to NSs, called speckle associated domains (SPADs) (27, 30, 32, 57). We asked if the observed defects in NS colocalization might correlate with changes in overall levels of HIV-1 integration as well as sites of integration within the human genome. To detect total levels of HIV-1 integration, we utilized Alu-Gag qPCR, which amplifies sequences that lay between the HIV-1 *gag* gene and genomic Alu-repeats; a second, nested qPCR was used to quantify the extent of first round reaction products (61, 62). We observed a significant reduction in integration in CPSF6-358 expressing cells, as expected, due to viral restriction in the cytoplasm (Fig 6A). Interestingly, we also observed a reduction in Alu-Gag products in cells expressing CPSF6-NP NLS and MX2 NLS (Fig 6A). CPSF6-SV40 NLS and C-MYC NLS by contrast supported Alu-Gag integration levels similarly as full-length CPSF6 (Fig 6A). These data are consistent with the notion that overall integration levels largely dictate levels of HIV-1 infection as assessed by viral reporter gene expression. To further probe the granularity of the integration phenotypes, we mapped sites of HIV-1 integration within the human genome essentially as described previously (63, 64). In short, asymmetric linkers were ligated onto sheared genomic DNA fragments, and LTR-linker sequences amplified via semi-nested PCR were analyzed by Illumina sequencing. Unique sites of HIV-1 integration were then mapped with respect to several genomic features, including genes, SPADs, lamina associated domains (LADs), and gene dense regions, as well as computationally-generated random integration control (RIC) values.

**Fig 6.**
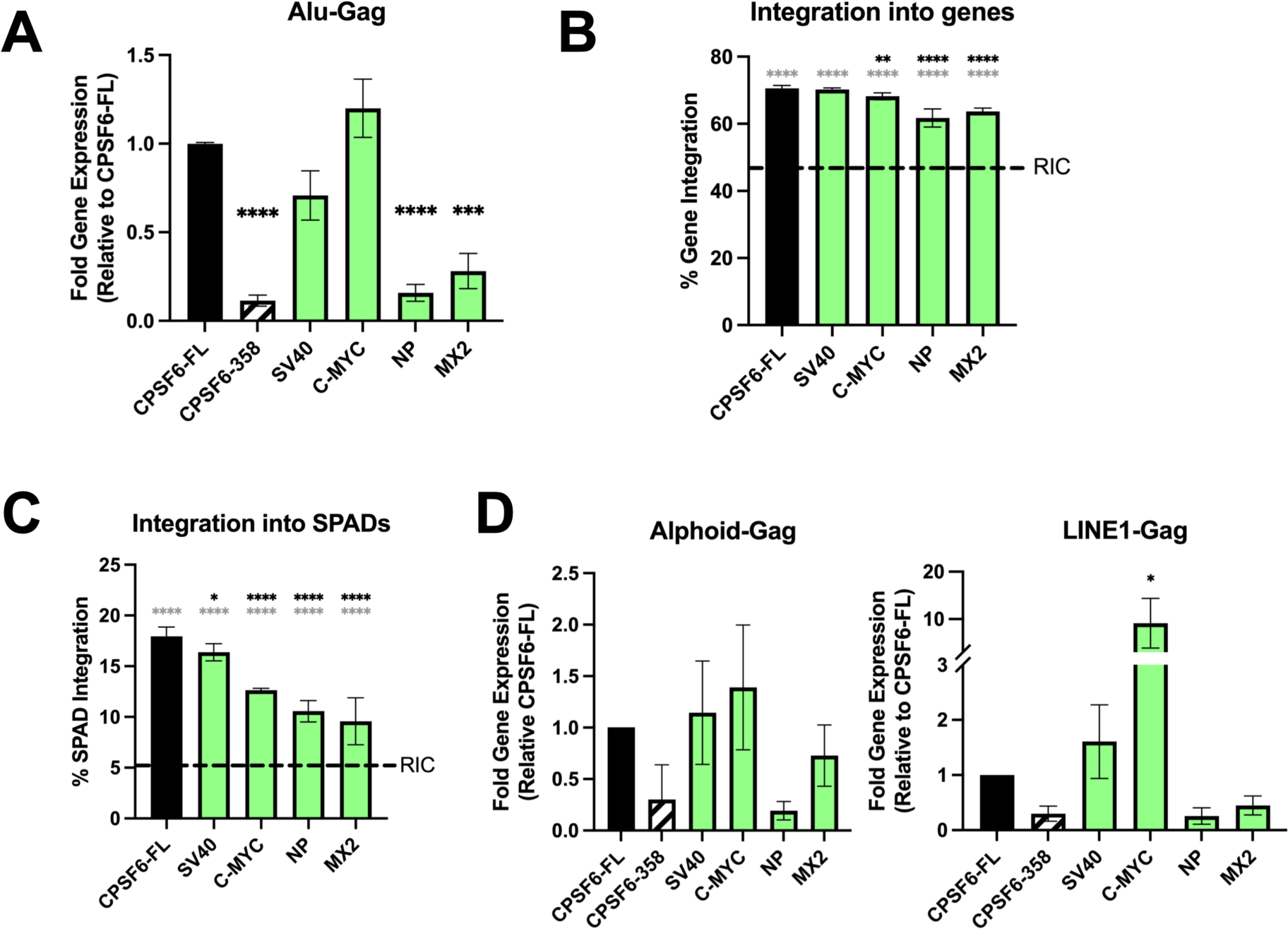
CPSF6-NP and MX2 NLS chimeras significantly impact HIV-1 integration. (A) Alu-Gag qPCR detection of HIV-1 integration levels. CPSF6-NP NLS and CPSF6-MX2 NLS cells supported significantly reduced levels of HIV-1 integration. (B) HIV-1 integration into genes. Integration into genes in CPSF6-C-MYC, NLS CPSF6-NP NLS, and CPSF6-MX2 NLS-expressing cells was reduced compared to cells expressing CPSF6-FL and CPSF6-SV40 NLS. (C) Integration into SPADs. P values in panels B and C compare results to CPSF6-FL expressing cells (black asterisks) and to indicated RIC values (grey asterisks). (D) Semi-nested Alphoid-Gag and LINE1-Gag qPCR was performed on genomic DNA from WT HIV-1 infected HeLa cells. Statistical analysis was determined using one-way ANOVA. Significant differences are indicated: * P<0.05, ** P<0.01, *** P<0.001, **** P<0.0001.

The number of unique integration sites mapped from a total of 2 to 4 independent infection experiments varied from 925 for the MX2 NLS chimera to 10,530 for C-MYC NLS protein (S1 Table). As expected, integration into genes in CPSF6-FL-expressing cells was significantly enriched compared to the RIC value (Fig 6B, grey asterisks). While genic integration in CPSF6-358 SV40 NLS-expressing cells was statistically indistinguishable from CPSF6-FL cells, ∼2-7% reductions in integration into genes was observed in C-MYC, NP, and MX2 NLS-expressing cells (Fig 6B and S1 Table). Overall similar trends were observed for integration into SPADs, LADs, and gene dense regions. Thus, while SPAD-proximal integration was marginally reduced by ∼1.7% in SV40 NLS-expressing cells (P<0.05), SPAD targeting was reduced by ∼5.4% to 8.7% in C-MYC, NP, and MX2 NLS-expressing cells (P<0.0001) (Fig 6C). Reciprocal upticks in integration into LADs in C-MYC, NP, and MX2 NLS-expressing cells differed significantly from LAD targeting values observed in CPSF6-FL and SV40-expressing cells (P<0.0001) (S2A Fig). In terms of gene dense regions, the 3.75 genes/Mb reduction in NP NLS-expressing cells was statistically significant versus the 15.4 genes/Mb value observed in CPSF6-FL cells. Interestingly, integration into gene dense regions in NP and MX2 NLS-expressing cells, compared to CPSF6-FL containing cells, was no longer enriched compared to the RIC 9.2 genes/Mb value (S2B Fig and S1 Table).

The human genome contains numerous types of repeated elements outside of Alu repeats, such as alphoid repeats that are enriched in centromeric regions and LINE1 repeats that are found throughout the genome. We used a modified Alu-Gag semi-nested qPCR approach that utilized primers specific for alphoid and LINE1 elements, as described previously (65). The CPSF6-MX2 NLS chimera facilitated integration into alphoid-enriched regions comparable to full-length CPSF6, whereas the CPSF6-NP NLS protein supported reduced levels of alphod repeat proximal integration (Fig 6D). While the SV40 NLS protein directed integration near LINE1 elements similar to CPSF6-FL, the CPSF6-C-MYC NLS chimera yielded a 10-fold increase in integration frequency near LINE1 elements (Fig 6D). As observed in the Alu-Gag assay, the NP and MX2 NLS proteins supported less LINE1-proximal integration than CPSF6-FL. Collectively, our results establish the importance of CPSF6’s NLS in facilitating the intranuclear events of HIV-1 infection.

## Discussion

In this study, we reveal that the NLS utilized by CPSF6 can impact various stages of nuclear trafficking and integration. By attaching heterologous NLSs to a truncated CPSF6, CPSF6-358, we observed a range of outcomes that were dependent on the NLS utilized. In line with previous experiments, some NLSs failed to complement the nuclear localization defect of CPSF6-358, and as a result restricted HIV-1 infection in the cytoplasm (39, 40). This is generally consistent with the ability of different nuclear import pathways, perhaps through heterogenous NPCs, to accommodate different nuclear transport cargoes (38, 39). We hypothesize that multiple NPC subsets can be found within a cell, and these NPC subsets have alternative nuclear transport mechanisms (39).

Following infection with HIV-1, we observed differences in reporter gene expression amongst the nuclear-rescued CPSF6-358 constructs. Some of the NLSs (C-MYC, Rac3) displayed a slight, but significant increase in reporter gene expression relative to full-length CPSF6. Additionally, the C-MYC NLS construct resulted in significantly increased integration as measured by Alu-Gag and LINE1-Gag qPCR assays. A possible explanation for this phenotype is that the C-MYC NLS pathway results in easier access to transcriptionally active chromatin, resulting in greater numbers of integrated provirus proximal to LINE1 elements and greater levels gene expression. Although omitted from downstream analyses, the Rac3 NLS protein behaved similar to the C-MYC protein in HIV-1 infection assays. Several additional constructs, including SV40, HNRNP K, and HNRNP A1, supported WT HIV-1 infection levels similar to CPSF6-FL.

Our results reveal that infection of CPSF6-NP and MX2 NLS cells resulted in functional nuclear import, as measured by both 2-LTR circles and fluorescently-tagged virus localization. However, we found that infection in these cell lines resulted in decreased levels of viral integration as measured by Alu-Gag and LINE1-Gag qPCR. The observation that 2-LTR circle formation was similar in cells with nuclear localized CPSF6 fusions suggests that the fusion constructs do not negatively impact core uncoating during infection, as 2-LTR circle formation requires the non-homologous end joining machinery. However, the diminished integration levels suggest that despite uncoating, the genome is unable to integrate into host chromatin. Our integration site analyses revealed that HIV-1 in CPSF6-NP and MX2 NLS-expressing cells generally disfavored regions associated with active chromatin, such as genes and SPADs, which could further reduce the levels of gene expression supported by these integrated proviruses. We speculate that the overall reduced levels of integration observed in CPSF6-NP and MX2 NLS cells combined with the propensity for these proviruses to disfavor active chromatin account for the comparatively poor expression profiles of these proviruses. Of note, our work fails to address why overall integration is significantly diminished in CPSF6-NP and MX2 NLS-expressing cells. Possibly, HIV-1 encounters a novel nuclear restriction factor(s) via the altered paths of nuclear import instilled by these NLSs. Further work will need to be done to explore this hypothesis.

Prior studies have investigated the differences in the nuclear import pathways of WT HIV-1 and the capsid mutants P90A (16, 37) and N74D (34), showing that binding to cyclophilin A/Nup153 and CPSF6, respectively, significantly impacted nuclear import pathway utilization. In our studies, the N74D CA mutant, which is defective for CPSF6 binding, predictably infected all CPSF6 chimeric construct expressing cells equally. However, infection with the P90A mutant resulted in differential infectivities relative to the WT virus. Specifically, P90A was less sensitive than WT HIV-1 to restriction in CPSF6-NP NLS cells, which was reproduceable following CsA treatment. Similarly, P90A displayed greater relative infectivity than WT HIV-1 in CPSF6-HNRNP K NLS cells, highlighting a potential role of cyclophilin binding influencing sensitivity in these cell lines. We speculate this may be due to P90A virus entering the nucleus via an alternative nuclear import pathway, as we and others have shown (16, 37, 38). In this case, perhaps the activity of the CPSF6-NLS effects in the nucleus is modulated by the nuclear import pathway utilized.

We also observed differential sensitivities to the CPSF6-NLS chimeras amongst related lentiviruses, as infection with HIV-2 and SIVmac highlighted further virus-specific phenotypes. Additionally, CPSF6-NLS chimeras displayed differential phenotypes, specifically in the SV40 and C-MYC NLS chimeras, in T cell and macrophage cell lines, where we observed SV40-NLS only partially rescued infection in T cells but fully rescued infection in macrophages and C-MYC-NLS rescued infection to WT levels in T cells but doubled infectivity in macrophages.

Previous reports have shown that the CPSF6 C-terminal RSLD is an IDR that can phase separate as a GFP-RSLD fusion protein in vitro (42). Moreover, purified mCherry-CPSF6 displays liquid-liquid phase separation activity in vitro (66). These observations raise the possibility that the RSLD plays a role in CPSF6 biomolecular condensate formation during HIV-1 infection (35, 41, 42, 58, 60). Our results would support this notion, as we found that only full-length CPSF6, which contains an intact RSLD, formed CPSF6 condensates following HIV-1 infection, while the CPSF6-358 constructs lacking an RSLD remained pan-nuclear in distribution. Interestingly, infection in cell lines expressing CPSF6-SV40 NLS and CPSF6-C-MYC NLS constructs retained HIV-1 NS colocalization. This was a surprising finding, suggesting that HIV-1 NS colocalization can occur independently of CPSF6 LLPS activity, as the constructs containing the SV40 and C-MYC NLS effectively trafficked HIV-1 to NSs despite lacking the RSLD.

Collectively, these results reveal that NLSs have roles beyond targeting cargo for nuclear import, such as impacting viral subnuclear localization following nuclear entry. To our knowledge, there is little precedent for post-nuclear entry roles for nuclear localization sequences. As such, our results utilizing HIV-1 infection as a readout for the nuclear function of NLSs provide a template to understand the nuclear trafficking or other activity that may be imparted by specific NLSs.

## Materials and Methods

### Cell Lines

THP-1, SupT1, and Jurkat cells were obtained from ATCC. 293T and HeLa cells were cultured in DMEM and THP-1, SupT1, and Jurkat cells were cultured in RPMI. Both mediums were supplemented with 10% tetracycline-depleted fetal calf serum (R&D Systems), 1000 U/ml of penicillin, and 1000 U/ml of streptomycin. THP-1 cells were differentiated into macrophages with 100 ng/ml phorbol 12-myristate 13-acetate (PMA) (Sigma-Aldrich) for 48 h, followed by 48 h rest with normal media without PMA before experiments.

### Constructs

Truncated CPSF6 containing amino acids 1-358 (CPSF6-358) from human isoform 1 was cloned into the pCW57.1 backbone (Addgene 71782). A hemagglutinin (HA) tag as well as heterologous NLSs (Table 1) were appended onto the C-terminus of CPSF6-358. These constructs were under a doxycycline inducible promoter, and expression was induced with 5 µg/ml Doxycycline for 48 h before viral infection.

### Virus and Vector Production

To generate viral vectors containing CPSF6-NLS constructs, 293T cells were seeded in 10 cm dishes and co-transfected with 4 µg pCW57-CPSF6-NLS, 4 µg psPAX packaging plasmid, and 2 µg pCMV-VSVg with polyethylenimine overnight. Media was changed the next morning and pseudotyped vector was collected 48 h post transfection. To generate pseudotyped HIV-1, HIV-2, SIVmac, N74D, and P90A viruses, 293T cells were seeded in 15 cm plates and transfected with 20 µg WT/N74D/P90A NL4.3-Luciferase-mCherry, HIV-2-Luciferase, or SIVmac-Luciferase (67) constructs with 5 µg pCMV-VSVg using polyethylenimine. The HIV-2-Luciferase construct was made by swapping the GFP gene of HIV-2-GFP for the firefly luciferase gene, as previously described for analogous equine infectious anemia constructs (32). Media was changed the following morning and virus was collected at 48 and 72 h post transfection. NL4.3-based plasmid pNLXLuc.R-U3-tag (64) was used to make virus for integration site distribution experiments. To generate Gag-integrase-Ruby viral particles, 293T cells were seeded in 10 cm plates and transfected with 4.5 µg R7ΔEnv, 3.5 µg Gag-integrase-Ruby, and 2 µg pCMV-VSVg using polyethylenimine. Viral particles were collected 48 h post transfection. All viruses and vectors were filtered through 0.45-µm filters (Merck Millipore). Virus was concentrated by spinning at 5000 g overnight at 4 °C. Reverse transcriptase (RT) units of viral preps were determined using SG-PERT assay as previously described (68). Viral infections were synchronized by spinoculation at 1200 g for 2 h at 13 °C.

### Stable Cell Line Generation

CPSF6-knockout cell lines were generated using pLentiCRISPRv2 with previously described guides (69) under blasticidin selection. For HeLa cells, a concentration of 5 µg/ml blasticidin was used and for THP-1, SupT1, and Jurkat a concentration of 2 µg/ml was used. Following HeLa cell selection, cells were sorted by flow cytometry and single-cell clonal populations were verified by western blot. CPSF6 knockout cells were transduced with viral vectors containing CPSF6-NLS constructs under puromycin selection. 5 µg/ml puromycin was used for selection of HeLa and 2 µg/ml puromycin was used for THP-1, SupT1 and Jurkat. Construct expression was verified by western blot after 48 h doxycycline induction.

### Antibodies and Chemicals

HIV-1 capsid, p24, was stained using mouse monoclonal anti-HIV-1 p24 from Santa Cruz Biotechnology (catalogue no. sc-69728). SC35 was stained using mouse monoclonal anti p-SC35 from Santa Cruz Biotechnology (catalogue no. sc-53518). GAPDH was stained with mouse monoclonal anti GAPDH from Santa Cruz Biotechnology (catalogue no. sc-47724). CPSF6 was stained using either CPSF6 polyclonal antibody from Invitrogen (catalogue no. PA5-41830) or from Abcam (catalogue no. ab175237). HA-tag was stained using either rabbit and HA-tag from Sigma-Aldrich (catalogue no. H6908) or mouse anti-HA-Tag (6E2) from Cell Signaling Technology (catalogue no. 2376). Secondary antibodies with conjugated fluorophores used in immunofluorescence experiments were purchased from Jackson Immunoresearch Laboratories. Aphidicolin (Cayman Chemicals, catalogue no. 14007) was used to growth arrest cells. Doxycycline hyclate (Sigma-Aldrich, catalogue no. D9891) was used to induce construct expression. CsA was purchased from Sigma-Aldrich. Hoechst was purchased from ImmunoChemistry Technologies (Catalogue no. 639).

### Western Blot

Cell lysates were prepared using NP-40 lysis buffer (100 mM Tris pH 8.0, 1% NP-40, 150mM NaCl) containing protease inhibitor cocktail (Roche) for 20 min with shaking on ice. Lysates were then centrifuged at 16000 g for 2 min and supernatant was collected. Protein concentration was determined using Pierce BCA Protein Assay Kit (Thermo Fisher Scientific). SDS was then added to lysates and incubated at 95 °C for 5 min. Equal amounts of total protein were loaded into a 4-15% gradient gel (Bio-Rad Laboratories). Proteins were then transferred to a nitrocellulose membrane (Bio-Rad Laboratories). Nitrocellulose membranes were incubated with specific primary antibodies overnight at 4 °C on a rocker. Secondary antibodies conjugated with horseradish peroxidase (Thermo Fisher Scientific) were added for 1 h at room temperature on a rocker. Chemiluminescence was detected using ProteinSimple imaging system (FluorChem E).

### Quantitative PCR (qPCR)

Cells were infected with equal RT Units as determined by SG-PERT assay. qPCR was used to determine late reverse transcription, 2-LTR circle formation, and integration. DNA was isolated from cells using DNeasy Blood and Tissue Kit protocol (QIAGEN). DNA concentration was determined using NanoDrop 2000 spectrophotometer and diluted to equal concentrations. 2-LTR Forward: 5’-AACTAGGGAACCCACTGCTTAAG-3’. 2-LTR Reverse: 5’-TCCACAGATCAAGGATATCTTGTC-3’ (70). Primers used to measure late reverse transcription were specific for the Firefly Luciferase gene present in our virus. Firefly Luc Fwd: 5’-CGGAAAGACGATGACGGAAA-3’. Firefly Luc Rev: 5’-CGGTACTTCGTCCACAAACA-3’. Human beta actin was used as housekeeping gene for normalization. HuBeta Actin Fwd: 5’-TCACCCACACTGTGCCCATCT-3’. HuBeta Actin Rev: 5’-CAGCGGAACCGCTCATTGCCAATGG-3’.

The Alu-gag PCR assay for HIV-1 integration was performed as described previously (62, 71, 72). Due to lentiviral transduction to generate cell lines, modifications were made to primer sets used. Lambda T sequence was added to the reverse primer of reaction 1 to add specificity and reduce background for reaction 2. Primers used for reaction 1 were as follows. Alu Forward: 5’-GCCTCCCAAAGTGCTGGGATTACAG-3’. LINE1 Forward: 5’-CTTCCAGTTTTTGCCCATTCAGT-3’. Alphoid Forward: 5’-GCAAGGGGATATGTGGACC-3’. Gag-LambdaT Reverse: 5’-AGTTTCGCTTACGTGGCATGTTCCTGCTATGTCACTTCC-3’. Second reaction primers for qPCR were as follows. HIV-1 Gag Forward: 5’-TCAGCCCAGAAGTAATAC-3’. Lambda T Reverse: 5’-AGTTTCGCTTACGTGGCAT-3’.

### Infectivity Assays

Cells were plated at equal densities and growth arrested in 5 µg/ml aphidicolin overnight. Infection was synchronized as described above. Infectivity was measured at 48 h post infection by either luciferase assay or flow cytometry. Following infection with luciferase reporter viruses, cells were lysed in Passive Lysis Buffer (Promega) and luciferase activity was measured using a luminometer (Promega, GloMax Navigator). With viruses harboring mCherry reporter system, mCherry positive cells were determined using flow cytometry (BD Bioscience, BD FACSCanto II cytometer).

### Microscopy

Images were acquired using a DeltaVision widefield fluorescence microscope (Applied Precision) with a digital camera (CoolSNAP HQ; Photometrics) and 1.4-numerical aperture 100X objective lens. Excitatory light was created by Insight SSI solid-state illumination module (Applied Precision). Images were acquired in several Z-stacks and deconvolution was performed using the softWoRx software v.7.0.0 (Applied Precision, GE Healthcare). Deconvolved images were analyzed using Imaris x64 v.7.6.4 (Bitplane). The surfaces or spots function of Imaris was used to analyze the signal of interest. The same algorithm was used for all images of an experiment. ImageJ was used to determine raw integrated density to quantify percentage of staining within the nucleus.

### Statistical Analysis

Prism 10.0 (GraphPad) was used to generate graphs and for statistical analysis. Statistical significance was determined using a one-way or two-way ANOVA analysis, where multiple comparisons were measured against full-length CPSF6 (CPSF6-FL). Data are graphed as mean ± standard error mean, except when indicated otherwise in figure legend. Kruskal-Wallis test was performed where indicated in figure legends. Significant differences are indicated: * P<0.05, ** P<0.01, *** P<0.001, **** P<0.0001.

### Integration Site Analysis

Infections were synchronously performed with U3-modified HIV-1 (64) to enable LTR-specific PCR amplification in cells containing pre-existing CPSF6-358 NLS lentiviral vectors at ∼1 MOI. Genomic DNA was isolated from infected cells 72 h post infection using Quick-DNA Miniprep kit (Zymo Research, catalogue no. D3024).

Integration site sequencing libraries were prepared essentially as described previously (63, 64). Briefly, 5 µg genomic DNA was digested overnight with a restriction enzyme cocktail containing 100 U each of NheI-HF, AvrII, SpeI-HF, and BamHI-HF (New England Biolabs). Digested DNA was purified using a GeneJet PCR purification kit.

Freshly annealed asymmetric linkers were subsequently ligated to the fragmented DNA in four parallel overnight reactions. These reaction products were then purified and subjected to two rounds of ligation mediated (LM-) PCR. The LM-PCR products were then purified and yield was quantified by DNA fluorimetry (Qubit). LM-PCR products were pooled at 10 nM total concentration, ΦX174 control DNA was added to the pooled libraries at 30% molar concentration, and the mixture was loaded onto a P2 300 cycle cartridge at 650 pM concentration for sequencing on an Illumina NextSeq2000 sequencer. Resulting raw FASTQ files were demultiplexed with Sabre (https://github.com/najoshi/sabre), the demultiplexed reads were trimmed to remove HIV-1 LTR and linker sequences, and the trimmed reads were aligned to human genome build hg19 as described (63). Uniquely mapped integration site annotations were used in downstream analyses. SPAD annotations used here were the same as previously described (27, 73). LAD annotations were downloaded from the 4D Nucleome data portal (Experiment Set 4DNESTAJJM3X). Integration site overlap with genomic features (genes, SPADs, and LADs) and gene density within 1 Mb of integration sites were quantified with the GenomicRanges Bioconductor package (74).

RIC datasets were generated by randomly placing integration sites within hg19 *in silico*. Following integration site placement, the genome was cleaved at the nearest downstream NheI, AvrII, SpeI, or BamHI recognition site. Generated fragments were filtered to be between 14 bp and 1200 bp and paired-end sequencing reads were simulated by taking up to 150 bp from both fragment ends to generate simulated R1 and R2 fasta files. These fasta files were then aligned to the hg19 genome and the location of uniquely mappable random integration sites were obtained. For statistical analysis of feature overlap, replicate datasets for each condition were pooled and Fisher’s exact test was used to test for equal proportions between conditions. Student’s t-test was used to assess statistical significance of gene density around integration sites between conditions.

## Data Availability Statement

Illumina Sequencing results can be found with Accession no: PRJNA1123763. All other relevant data are within the paper and its Supporting Information files.

## Supporting Information

**S1 Fig.**
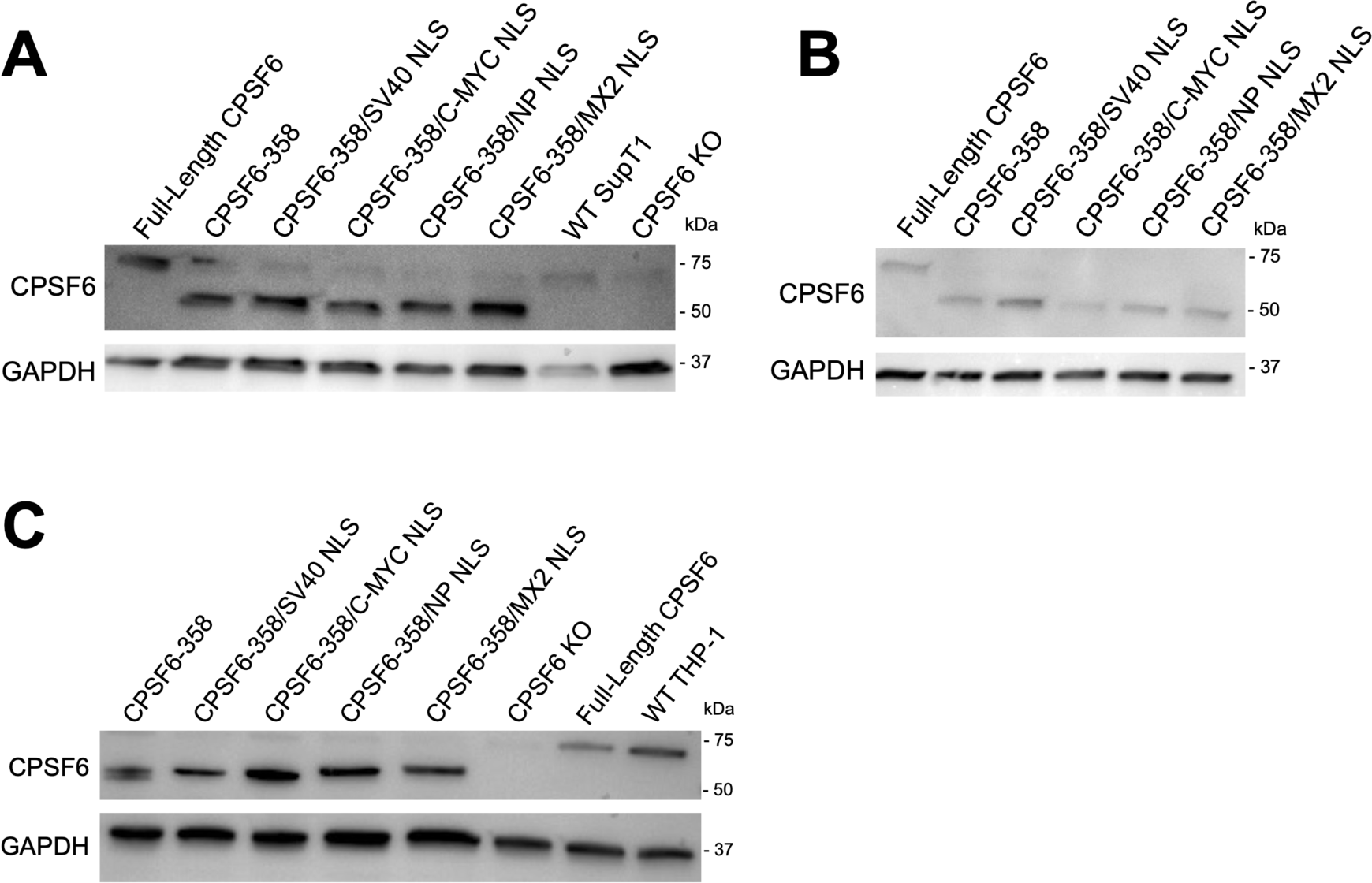
Chimeric construct expression in T cells and macrophages. Western blot analysis of CPSF6-depleted stably-transduced SupT1 (A), Jurkat (B), and THP-1 (C) cell lines using anti-CPSF6 antibody to detect CPSF6-NLS construct expression following 48 h doxycycline induction. Anti-GAPDH antibody used as loading control.

**S2 Fig.**
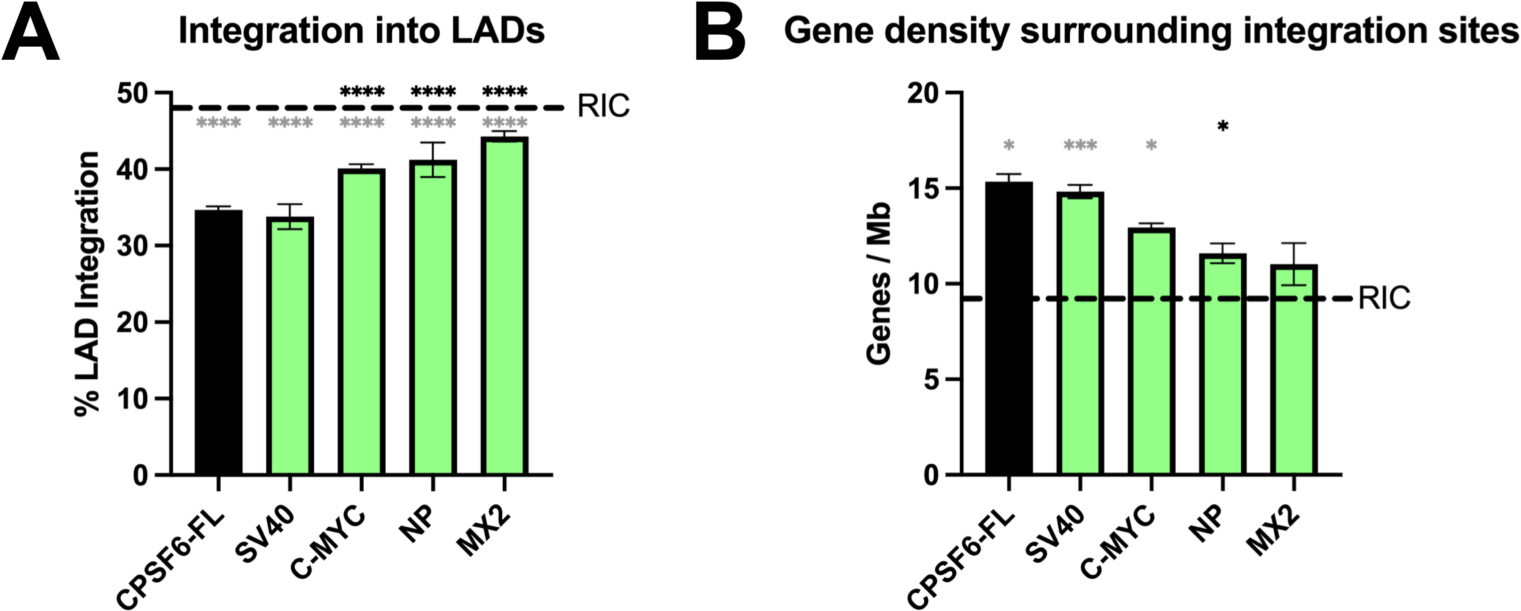
Integration site targeting metrics. (A) HIV-1 integration targeting with respect to LADs for the indicated cell lines. Fractional LAD targeting in CPSF6-C-MYC, NLS CPSF6-NP NLS, and CPSF6-MX2 NLS-expressing cells was increased significantly compared to cells expressing CPSF6-FL and CPSF6-SV40 NLS. (B) Integration into gene rich regions of chromatin. Integration in gene dense regions in CPSF6-NP NLS cells was reduced significantly compared to CPSF6-FL expressing cells. Integration in NLS CPSF6-NP NLS and CPSF6-MX2 NLS cells was interestingly statistically indistinguishable from the RIC value. * P<0.05, *** P<0.001, **** P<0.0001 (black asterisks, vs CPSF6-FL cells; grey asterisks, vs corresponding RIC).

**S1 Table.** Integration distributions in CPSF6-NLS chimera HeLa cells^a^.

| Sample<sup>b</sup> | Unique Sites | % Integration in Genes | % LAD Integration | % SPAD Integration | Gene Density Surrounding Integration Site |
| --- | --- | --- | --- | --- | --- |
| C6FL Ctrl | 59 |  |  |  |  |
| C6FL inf A | 3007 | 69.74 | 34.25 | 512 | 14.94 |
| C6FL inf B | 3951 | 71.40 | 35.16 | 745 | 15.74 |
| SV40 ctrl | 28 |  |  |  |  |
| SV40 inf A | 2236 | 70.71 | 32.16 | 385 | 15.18 |
| SV40 inf B | 1667 | 69.71 | 35.45 | 259 | 14.48 |
| C-MYC Ctrl | 42 |  |  |  |  |
| C-MYC inf A | 5864 | 69.24 | 39.56 | 752 | 13.16 |
| C-MYC inf B | 4666 | 67.19 | 40.63 | 581 | 12.72 |
| NP Ctrl | 18 |  |  |  |  |
| NP inf A | 315 | 59.05 | 43.49 | 30 | 11.08 |
| NP inf B | 844 | 64.45 | 38.98 | 98 | 12.10 |
| MX2 Ctrl | 21 |  |  |  |  |
| MX2 inf A | 429 | 62.70 | 43.59 | 51 | 12.12 |
| *MX2 inf B* | 496 | 64.72 | 44.96 | 36 | 9.92 |
| *C6KO Ctrl* | 87 |  |  |  |  |
| *C6KO inf* | 2273 | 62.74 | 44.13 | 220 | 11.50 |
| *RIC* | 112183 | 45.09 | 51.20 | 3178 | 7.88 |

<sup>a</sup>Results are from two independent infection experiments (inf A and inf B).
<sup>b</sup>C6FL – full-length CPSF6; C6KO – CPSF6-knockout; RIC – random integration control.

